# Evidence for a role of extraintestinal pathogenic *Escherichia coli, Enterococcus faecalis* and *Streptococcus gallolyticus* in the aetiology of exudative cloacitis in the critically endangered kākāpō (*Strigops habroptilus*)

**DOI:** 10.1101/2024.11.07.622535

**Authors:** Rebecca K. French, Stephanie J. Waller, Janelle R. Wierenga, Rebecca M. Grimwood, James Hodgkinson-Bean, Andrew Digby, Lydia Uddstrom, Daryl Eason, Kākāpō Recovery Team, Lisa S. Argilla, Patrick J. Biggs, Adrian Cookson, Nigel P. French, Jemma L. Geoghegan

**Affiliations:** Department of Microbiology and Immunology, University of Otago, Dunedin, New Zealand; Kākāpō Recovery Programme, Department of Conservation/Te Papa Atawhai, Invercargill, New Zealand; Dunedin Wildlife Hospital, Otago Polytechnic School of Animal Health, New Zealand; AgResearch, Hopkirk Research Institute, Palmerston North, New Zealand; mEpiLab, School of Veterinary Sciences, Massey University, Palmerston North, New Zealand; Tāwharau Ora | School of Veterinary Science, Massey University, Palmerston North, New Zealand

## Abstract

The kākāpō is a critically endangered flightless parrot which suffers from exudative cloacitis, a debilitating disease resulting in inflammation of the vent margin or cloaca. Despite this disease emerging over 20 years ago, the cause of exudative cloacitis remains elusive. We used total RNA sequencing and metatranscriptomic analysis to characterise the infectome of lesions and cloacal swabs from nine kākāpō affected with exudative cloacitis, and compared this to cloacal swabs from 45 non-diseased kākāpō. We identified three bacterial species – *Streptococcus gallolyticus*, *Enterococcus faecalis* and *Escherichia coli* – as significantly more abundant in diseased kākāpō compared to healthy individuals. The genetic diversity observed in both *S. gallolyticus* and *E. faecalis* among diseased kākāpō suggests that these bacteria originate from exogenous sources rather than from kākāpō-to-kākāpō transmission. The presence of extraintestinal pathogenic *E. coli* (ExPEC)-associated virulence factors in the diseased kākāpō population suggests that *E. coli* may play a critical role in disease progression by facilitating iron acquisition and causing DNA damage in host cells, possibly in association with *E. faecalis*. No avian viral, fungal nor other parasitic species were identified. These results, combined with the consistent presence of one *E. coli gnd* sequence type across multiple diseased birds, suggests that this species may be the primary cause of exudative cloacitis. These findings shed light on possible causative agents of exudative cloacitis, and offer insights into the interplay of microbial factors influencing the disease.

## 1. Introduction

Accurately diagnosing disease in critically endangered and evolutionary divergent species is challenging because of the inability to perform challenge studies coupled with the lack of model host organisms or cell lines. In addition, many molecular diagnostic techniques are specific to certain microorganisms, which can hamper disease investigation. For example, 16S or 18S rRNA sequencing will only detect bacteria or eukaryotic parasites, respectively, but not viruses, and polymerase chain reaction (PCR) may be too specific to detect novel organisms. Total RNA sequencing, or metatranscriptomics, on the other hand, can identify the entire ‘infectome’ including viruses, bacteria, fungi and other eukaryotic parasites, as well as the expression of genes, including those involved in virulence and antimicrobial resistance (Van Borm et al., 2015). Therefore, metatranscriptomics is an ideal tool for pathogen investigation, particularly in critically endangered species.

The kākāpō is a critically endangered flightless parrot endemic to Aotearoa New Zealand and the only extant member of the genus *Strigops*. Once widespread across the archipelago, following severe population decline (Bergner et al., 2016) the species was considered possibly extinct by the 1960s before being rediscovered in remote Fiordland and Rakiura/Stewart Island in the 1970s (Clout & Merton, 1998). Since their rediscovery, extensive conservation efforts (Elliott et al., 2001) have resulted in a gradual population increase, with a current population size of 244 across six islands (Department of Conservation). The extreme population bottlenecks have resulted in a high level of inbreeding which may increase the species’ vulnerability to disease, although recent research suggests the population does not have a higher deleterious mutation load (Dussex et al., 2021; Guhlin et al., 2023). In any case, given that their population size is still extremely small, the kākāpō is highly vulnerable to stochastic events such as pathogen emergence and disease outbreaks.

In 2003, a disease termed exudative cloacitis emerged in kākāpō (Jakob-Hoff et al., 2009). Exudative cloacitis is a debilitating disease resulting in inflammation of the vent margin or cloaca, which often becomes ulcerated and covered in crusty exudate. As kākāpō are wild animals (although highly managed), the disease is usually diagnosed during routine health checks, or from a decline in activity detected by remote monitoring of activity transmitters. Kākāpō with mild cases are usually left to recover naturally, while more severe cases are removed from the island and treated in a wildlife hospital with antibiotics (primarily Amoxicillin/clavulanic acid). Until recently this disease was confined to one of the three island populations (Whenua Hou/Codfish Island, Figure 1), but in 2021 the disease spread to a second island (Te Kākahu/Chalky Island) and to a third island in 2022 (Pukenui/Anchor Island) (unpublished data). This occurred in spite of strict quarantine procedures in place for transfer of people and equipment between the islands, which includes disinfection of clothing and equipment. Outbreaks occur sporadically and the number of cases per year is variable, although appears to be climbing (unpublished data). In addition, as the kākāpō population increases, the ability to identify and individually treat diseased birds will become less practical. Thus, given the increase in cases and widening geographic spread, this disease is becoming an increasing threat to kākāpō conservation.

**Figure 1.**
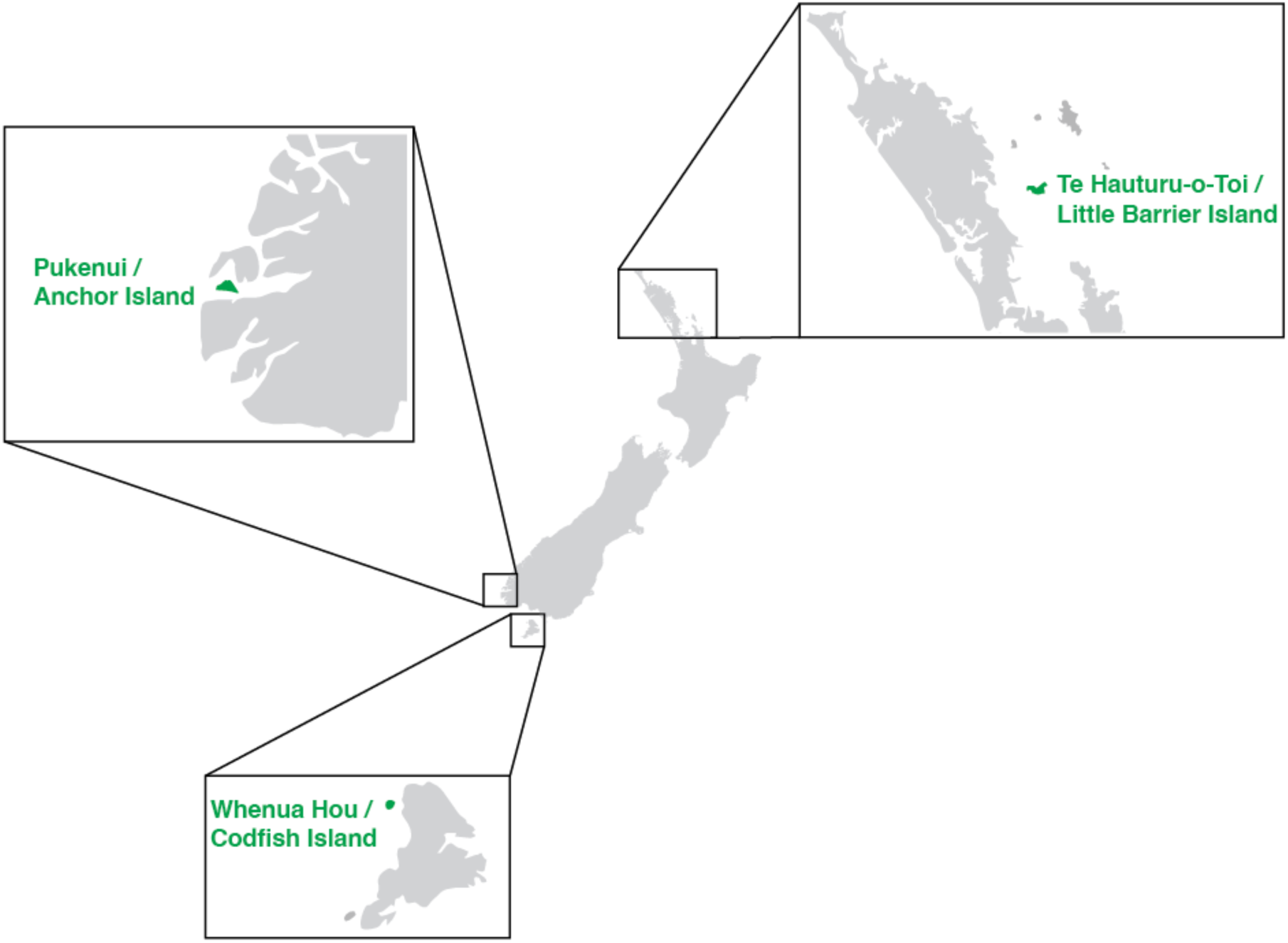
Map of New Zealand showing the three islands that kākāpō were sampled from (green).

Despite this disease emerging over 20 years ago, the cause of exudative cloacitis remains unknown. Diagnostic testing including culture of lesions and faeces, haematology, blood biochemistry and PCR for *Psittacine circovirus* of one affected individual failed to determine the cause (Jacob-hoff & Gartrell, 2011). Metagenomics of another single case (compared to a pool of eight healthy individuals) tentatively linked Escherichia phage TL-2011b to the disease, but given the small sample size this was considered only a ‘plausible hypothesis’ (White et al., 2015), and subsequent PCR-based testing did not support these findings (unpublished data). An investigation into possible environmental factors did not find any strong evidence to link these factors to the disease (Jayasinghe et al., 2022). A metagenomic/metatranscriptomic study of primarily faecal samples from nine diseased and three healthy kākāpō was unable to identify a potential causative agent for exudative cloacitis (West et al., 2024). However, the study did identify differential expression of some bacterial genera including *Enterococcus*, *Escherichia*-*Shigella*, *Streptococcus*, as well as potential links to the expression of toxin producing *hok*/*gef* genes (West et al., 2024), which compel further investigation. To date, there have been no metagenomic or metatranscriptomic investigations of diseased lesion tissue, which may provide better insight into potential causative pathogens.

Herein, we characterised the infectome of lesions and cloacal swabs from nine kākāpō affected with exudative cloacitis, and compared this to cloacal swabs from 45 non-diseased kākāpō in order to elucidate the possible cause of exudative cloacitis.

## 2. Methods

### 2.1 Sampling

Cloacal swabs were taken by the Department of Conservation from wild kākāpō caught for routine health checks from three offshore predator-free islands – Te Hauturu-o-Toi/Little Barrier Island (n = 3), Pukenui/Anchor Island (n = 4) and Whenua Hou/Codfish Island (n = 54) (Figure 1). These checks are usually annual, although may be more frequent during breeding periods, or if a kākāpō shows unusual activity patterns from remote monitoring of activity transmitters. Kākāpō that were healthy at the time of sample collection but had a previous exudative cloacitis infection were included along with their infection history. Kākāpō displaying active exudative cloacitis symptoms were either sampled in the wild when caught for health checks, and/or, for those transferred for treatment, in the Dunedin Wildlife Hospital. Where lesions were present, tissue of the lesions were sampled, and a swab taken from the lesion and cloaca. Tissue samples and swabs were placed in 800µl of DNA/RNA Shield (Zymo Research) or RNAlater stabilization solution (ThermoFisher), and kept at -20°C before transfer to a - 80°C freezer.

### 2.2 RNA extraction and sequencing

Defrosted cloacal swabs or lesions were placed in ZR BashingBead Lysis Tubes (0.1 mm and 0.5 mm) (Zymo Research) filled with 1 mL of DNA/RNA shield (Zymo Research).

Lysis tubes were homogenized for five minutes in a mini-beadbeater 24 disruptor (Biospec Products Inc.). RNA was extracted using the ZymoBIOMICS MagBead RNA kit (Zymo Research) as per the manufacture’s protocol. In brief, RNA was extracted using magnetic bead-based purification, and eluted into 50μl DNase/RNase free water.

Extracted RNA from diseased kākāpō (n = 9) was kept separate (i.e. only samples from the same kākāpō were pooled together), including from cloacal swabs and sampled lesions (Figure 2). RNA from healthy individuals was pooled in equal volumes according to sample type and island, as well as past infection status, to increase RNA quantity, resulting in a total of 34 RNA pools (Figure 2). Complementary DNA libraries were prepared using the Illumina Stranded Total RNA Prep with Ribo-Zero Plus. RNA was sequenced on the Illumina NovaSeq 6000 platform at AgResearch (Invermay), Otago, New Zealand.

**Figure 2.**
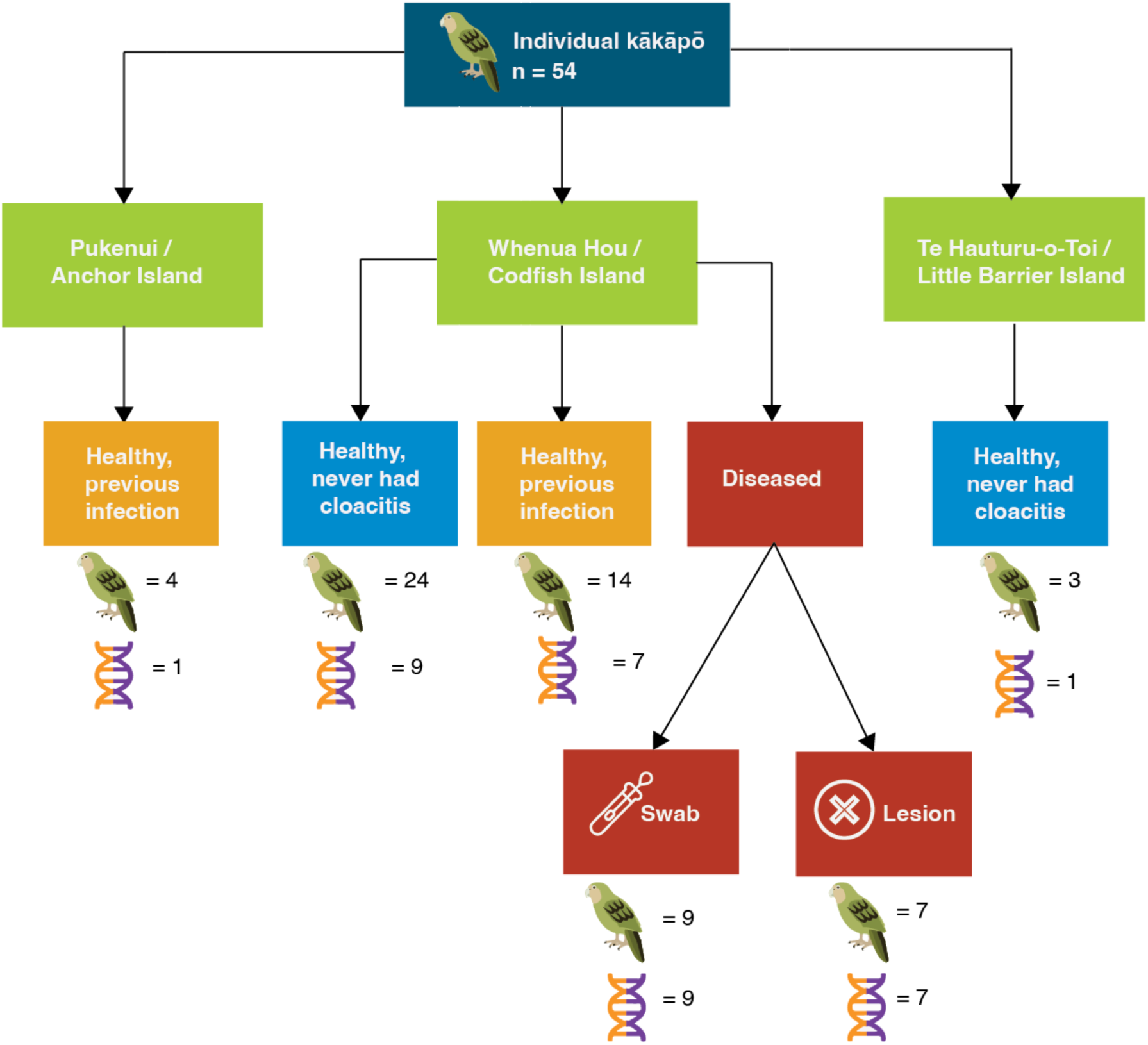
Map showing pooling strategy. Kākāpō samples were pooled according to island, disease status and sample type. All samples from healthy birds were swabs. The kākāpō icon denotes the number of individual kākāpō sampled, the helix icon denotes the number of RNA libraries. RNA from individual diseased kākāpō was pooled.

### 2.3 Bioinformatic pipeline

Trimmomatic (v0.38) was used to trim sequencing adapters and bases with a quality score below five, applying a sliding window with a window size of four (Bolger et al., 2014). Additionally, bases with a quality score below three at the start and end of the reads were removed. Sequences shorter than 100 nucleotides or with an average quality below 10 were removed using bbduk in BBtools (bbmap v39.01, (Bushnell, 2014). *De novo* assembly of the reads into longer contiguous sequences (contigs) was performed using MEGAHIT (v1.2.9) (Li et al., 2015). Bowtie2 (v2.4.5, (Langmead & Salzberg, 2013)) and SAMtools (v1.9, (Li et al., 2009)) were used to estimate the abundance of each contig.

### 2.4 Virus discovery and phylogenetic analysis

Viral contigs were identified by comparing the assembled sequences to the National Center for Biotechnology Information (NCBI) nucleotide (nt) and non-redundant protein (nr) databases using BLASTN (BLAST+ v2.13.0, (Camacho et al., 2009)) and DIAMOND BLASTX (DIAMOND v2.1.6, (Buchfink et al., 2015)). To reduce false positives, sequence similarity cut-off values of 1 × 10^−5^ for nt and 1 × 10^−10^ for nr were applied.

A protein structure similarity-based search was used to identify highly divergent viral transcripts that did not share sequence similarity to other known transcripts within NCBI nt and nr databases, termed orphan contigs. Orphan contigs were first translated into open reading frames (ORFs) using the EMBOSS getorf program (Rice et al., 2000), with the minimum nucleotide size set to 1,000 nucleotides, the maximum nucleotide size set to 50,000 and the “methionine” flag option set to only report ORFs with a methionine amino acid starting codon. Translated ORFs were then used as queries against the RdRp Scan HMM profile database (Charon et al., 2022) to identify putative viral polymerase sequences. ColabFold (Mirdita et al., 2022) was then used to model putative viral polymerase sequence structures and Foldseek (Van Kempen et al., 2024) was used to align putative viral polymerase structures against the PDB100 database (Velankar et al., 2021) using the 3Di/AA mode.

To identify other highly divergent viruses we screened the protein sequence identified via the protein structure similarity approach against transcriptome assemblies available in NCBI’s Transcriptome Shotgun Assembly (TSA) sequence database (Forgia et al., 2022) using the translated BLAST tool. Searches were restricted to eukaryotes (taxid:2759). Putative viral sequences were analyzed using Geneious Prime 2022.2.2 to find and translate ORFs. Translated protein sequences were then queried against the online nr protein database (database accessed June 2024) using the BLASTp tool.

Translated sequences of the putative viral polymerase structures were aligned with representative protein sequences from the same taxonomic viral clade obtained from Forgia et al. (2022) and the translated TSA hits using MAFFT v7.450 (L-INS-I algorithm). Poorly aligned regions were removed using trimAL v1.2 (Capella-Gutiérrez et al., 2009) with the gap threshold flag set to 0.9. IQ-TREE v1.6.12 was used to estimate maximum likelihood phylogenetic trees and the LG amino acid substitution model was selected with 1000 ultra-fast bootstrapping replicates (Nguyen et al., 2015).

### 2.5 Identification of non-viral microorganisms

The microbial communities (including bacteria, fungi, archaea and eukaryotic parasites) in each sample were characterized using CCmetagen (v1.4.3, (Marcelino et al., 2020)), with an abundance cut-off of one read per million (RPM). An operational taxonomic unit table was created using abundance expressed as the number of reads per million (RPM, that is, the number of reads from each microbial family divided by the total number of reads in the library, multiplied by one million). Alpha diversity (richness – number of microbial families and abundance – reads per million) was calculated, and the role of health status and sample type (swab and lesion) were interrogated using generalized linear mixed effect models with a poisson link (richness, count data) and general linear mixed effect models (abundance) using R (v4.3.1) and the lme4 R package (v1.1-34, (Bates et al., 2015)). In all models, kākāpō identity was used as a random effect to account for there being multiple libraries from the same diseased individual. While the libraries from healthy birds were pooled, individual birds were always in separate libraries, and so each healthy library was treated as a separate individual. Beta diversity (that is, shared diversity across libraries) was visualized at the microbial family level using principal coordinate analysis (PCoA) with a Bray–Curtis dissimilarity matrix, presented as a PCoA plot using the R packages phyloseq v1.44.0 (McMurdie & Holmes, 2013) and vegan v2.6-4 (Oksanen et al., 2013). Bray–Curtis dissimilarity is a statistic ranging from 0 to 1 that reflects the dissimilarity of communities between libraries, with 0 meaning both libraries have an identical community, and 1 meaning the libraries have no microbial families in common. Pairwise permutational analyses of variance (PERMANOVA, adonis2 in the vegan R package) was used to test for differences in the community based on health status and sample type, using the Bray–Curtis dissimilarity matrix and alpha of 0.05.

Differential expression analysis of the abundances of each genus (including all bacterial, fungal and eukaryotic genera, excluding host RNA) was conducted using the R package DESeq2 (v1.40.2, (Love et al., 2014)). Genera were considered potential pathogens and analyzed further if they met the following criteria: (1) the genus was present in over half of the diseased libraries, and (2) the genus was present in a significantly higher abundance (using an adjusted alpha of 0.05 following a Bonferroni correction to correct for multiple comparisons and control for potential Type I error) in the diseased compared to the healthy libraries in the differential expression analysis.

Four bacterial genera met these criteria: *Streptococcus, Enterococcus, Corynebacterium* and *Escherichia.* Within these four genera, forty-nine bacterial species were represented in our dataset. The abundances of each of these species were compared using stacked bar plots (created in R using ggplot2), and the dominant species within this subset (*Streptococcus gallolyticus*, *Enterococcus faecalis*, and *Escherichia coli)* subject to further analysis, as described below.

For the most part, full seven-gene multilocus sequence typing (MLST) was not possible with our short-read, total RNA sequencing data as the individual alleles for each gene could not be matched together. Instead, the abundance and phylogeny of each locus from the MLST scheme was analyzed individually. For *S. gallolyticus* and *E. faecalis,* we used all seven loci from their respective MLST schemes and compared these to the PubMLST database (Jolley et al., 2018). For *E. coli*, we used the *gnd* (6- phosphogluconate dehydrogenase) locus for sequence typing (Cookson et al., 2017), and then conducted phylogenetic analysis using the entire *gnd* operon. Gene loci were extracted using an R based pipeline (available here: https://github.com/dw974/dnatools/blob/master/R/dnatools.R) involving a customized BLASTN search (BLAST+ 2.15.0) and MAFFT (v7.526) alignment, which then assigns an allele number to each sequence. Maximum likelihood phylogenetic trees of each locus were created in IQ-TREE v1.6.12 using these alignments with the best-fit substitution model determined by the ModelFinder (Kalyaanamoorthy et al., 2017) and node robustness assessed by employing the approximate likelihood ratio test with 1000 replicates (Nguyen et al., 2015).

A partial *E. coli* genome was created by mapping the assembled contigs to a reference whole genome CP028770 in Geneious (v2023.2.1 https://www.geneious.com), using the ‘map to reference’ tool. The SerotypeFinder CGE tool (Joensen et al., 2015) was used for *in silico* serotyping, and mlst 2.19.0 (Seemann T, mlst Github https://github.com/tseemann/mlst) was used for sequence type identification using the Achtman seven-locus multilocus sequence typing (MLST) scheme and PubMLST database (Jolley et al., 2018). A separate analysis was also conducted where assembled contigs were binned using metabat2 v2.15 (Kang et al., 2019) based on tetranuc (kmer=4) frequency and coverage, and MaxBin2 v2.2.6 (Wu et al., 2016). Bins were identified using DAS Tool v1.1.1 (Sieber et al., 2018) and Kraken2 v2.1.2 (Wood et al., 2019), resulting in two *E. coli* bins. These bins were identified in the same way as above (SerotypeFinder CGE tool and mlst), producing the same *in silico* serotyping and sequence type.

### 2.6 Virulence factors

Virulence factors of each bacterial species were identified using Bowtie2 (v2.4.5) and the R package Rsubread (v2.14.2) (Liao et al., 2019) to map the trimmed reads to annotated whole genomes and quantify the read abundance for each known virulence gene (based on the Virulence Factors of Pathogenic Bacteria database (Chen et al., 2005), and published literature (Danne et al., 2011; de Oliveira Alves et al., 2024; Johnson et al., 2008; Klemm et al., 2010; Lüthje & Brauner, 2010; Nilsson et al., 2016; Shankar et al., 2006; Stępień-Pyśniak et al., 2019). Differential expression analysis of the abundances of each gene was conducted using the R package DESeq2 (v1.40.2).

### 2.7 Host resistome

The KMA (k-mer alignment) program (v1.4.15, (Clausen et al., 2018)) was used to map trimmed reads to the ResFinder (Bortolaia et al., 2020) and PointFinder (Zankari et al., 2017) databases to identify antimicrobial resistance (AMR) genes. This analysis was run for all AMR genes from the entire ResFinder database, and separately from the *E. coli* and *E. faecalis* PointFinder databases. The ResFinder database contains AMR genes, while the PointFinder database contains chromosomal point mutations mediating resistance for specific bacterial species. Both databases are compiled from published literature and pre-existing databases, and curated by experts (Bortolaia et al., 2020).

The databases are regularly updated and were downloaded in August 2024. As described above, alpha diversity (richness and abundance) was calculated, and the role of health status and sample type (swab and lesion) on the abundance (expressed as RPM) of all and individual AMR genes or point mutations were interrogated using linear mixed effects models.

## 3. Results

### 3.1 The kākāpō metatranscriptome

The kākāpō metatranscriptome was sequenced at an average of 80.9 million reads per library (range 31 million – 157 million). There was no significant difference in RNA sequencing depth according to health status (p = 0.48, Supplementary Table 1) or sample type (p = 0.15, Supplementary Table 1). On average, 11% (range 8 – 19%) of the assembled contigs were orphan contigs (i.e. they had no significant sequence similarity in reference databases). The overall metatranscriptome across all sample types consisted primarily of bacteria (74%), followed by eukaryotes (25%), viruses (0.09%) and archaea (0.07%).

### 3.2 Viruses in kākāpō

We found a significantly higher number of viral families in healthy libraries compared to diseased libraries (p = 0.008, Supplementary Table 1), and a significantly higher number of viral families in lesions compared to swabs (p = 0.04, Supplementary Table 1).

However, we found no differences in abundance according to disease status (p = 0.2, Supplementary Table 1) or sample type (p = 0.07, Supplementary Table 1). A principal co-ordinate analysis plot based on the Bray-Curtis dissimilarity matrix revealed weak clustering according to health status but not sample type, which was confirmed with a PERMANOVA test (p = 0.01 and p = 0.6 respectively, Supplementary Table 1).

We found viruses from 74 different viral families/unclassified groups. However, most of these viral taxa were present at a very low abundance, with the majority (51%) having a total abundance (i.e. the sum across all 34 libraries) of <10 RPM. ‘Unclassified viruses’ (i.e. those with a closest genetic relative to viruses classed as ‘unclassified viruses’ by NCBI taxonomy) were the most abundant overall (17,579 RPM), and appeared to be bacteriophages based on their other known genetic relatives. This was followed by *Caudoviricetes* (10,588 RPM, also bacteriophages), *Reovirales* (8,313 RPM, mite- infecting viruses) and unclassified *Riboviria* (1,249 RPM) that appeared to be fungi/plant infecting viruses. ‘Unclassified viruses’ and *Caudoviricetes* were present in all libraries.

For viral families that could be classified to family level, there were 28 families with a total abundance of >10 RPM. The majority of these were plant-infecting (54%), followed by bacteriophage (21%). We found no viruses that appeared to be infecting the kākāpō host (i.e. avian viruses) based on their sequence similarity.

A structural homology-based approach was also used to identify highly divergent viral sequences that could not be detected using sequence similarity-based approaches. Using this approach we identified one highly divergent virus that shared structural similarity to *La Crosse virus* (*Peribunyaviridae*) polymerase sequence (PDB 7ORO) with a probability 1, sequence identity 9.8% and e-value 5.59x10^-8^. We screened this highly divergent protein sequence against the TSA database and identified eight highly divergent ormycoviruses (including *Stalked jellyfish associated ormycovirus*, *Antarctic polychaete worm associated ormycovirus*, *Bean rust fungi associated ormycovirus*, *Nematode associated ormycovirus*, *Squid associated ormycovirus*, *Termite associated ormycovirus 1*, *Protozoan associated ormycovirus* and *Termite associated ormycovirus 2*). Upon phylogenetic analysis, it was clear that this virus fell within the ormycovirus clade and was likely infecting fungi rather than kākāpō (Supplementary Figure 1).

### 3.3 Microorganisms associated with disease in kākāpō

The overall abundance of all microorganisms (i.e. the infectome) was not significantly different between healthy and diseased kākāpō (Figure 3A, Figure 3D, p>0.1, Supplementary Table 1). However, this was not the case for sample type, with lesions having a significantly higher microbial abundance than swabs (Figure 3A, Figure 3D, p = 0.004, Supplementary Table 1). Libraries comprising healthy individuals had significantly higher microbial diversity (i.e. number of taxonomic families) than diseased birds (Figure 3B, Figure 3D, p = 0.002, Supplementary Table 1). A principal co-ordinate analysis plot based on the Bray-Curtis dissimilarity matrix revealed clustering according to health status and sample type (Figure 3C), which was confirmed with a PERMANOVA test (Figure 3D, p = 0.0001 and p = 0.007, respectively – Supplementary Table 1). Healthy kākāpō clustered tightly together indicating they share a similar microbial community, while diseased kākāpō had a greater variation with little to no clustering (Figure 3C), suggesting that multiple pathogens are potentially associated with exudative cloacitis.

**Figure 3.**
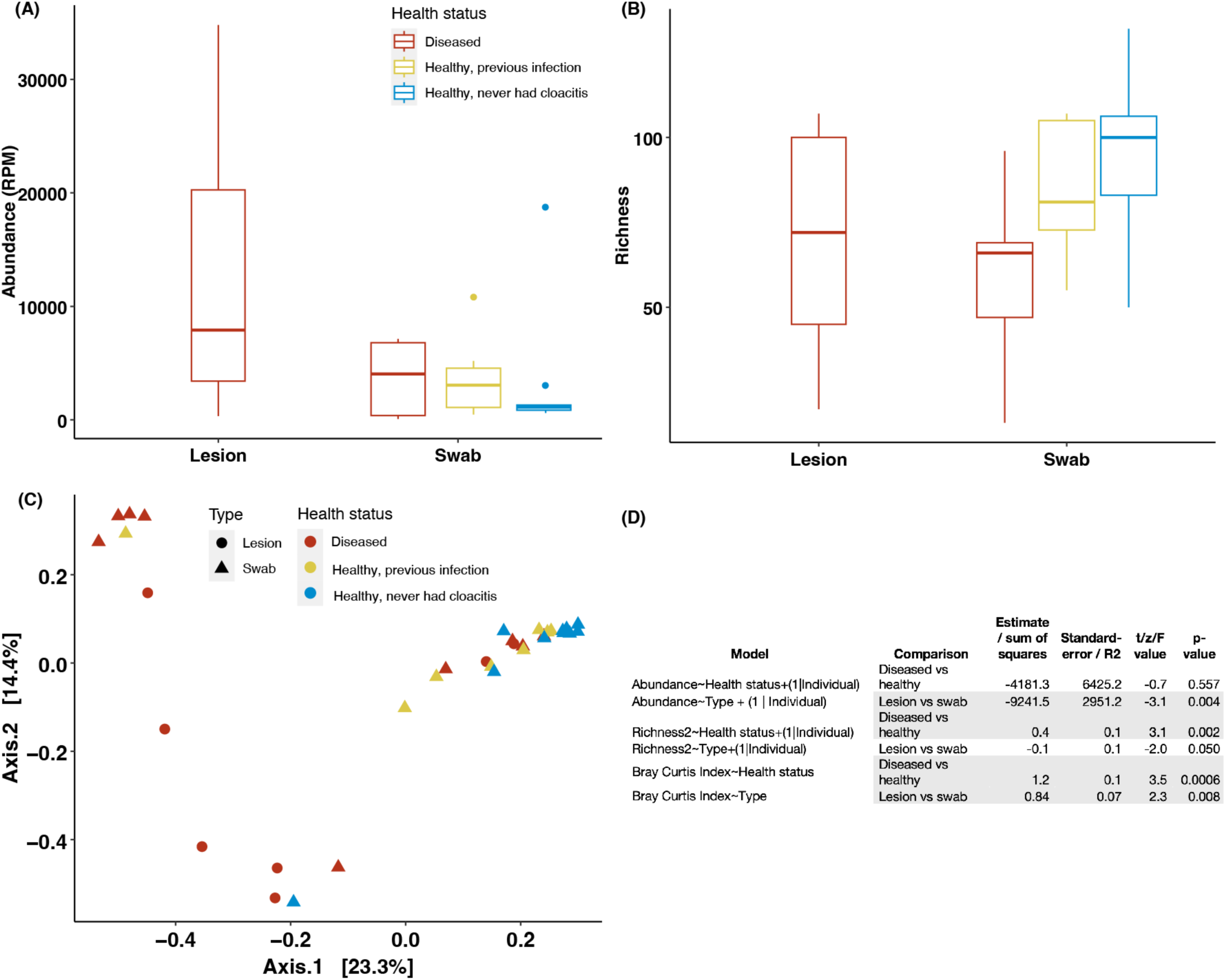
(A) Boxplot showing microbial abundance (reads per million) for lesions and swabs of differing health status. (B) Boxplot showing microbial richness (number of families) for lesions and swabs from kākāpō of differing health status. (C) The first two dimensions of a principal coordinate analysis (PCoA) plot showing microbial community clustering according to health status and sample type. (D) Linewar mixed effects model and PERMANOVA analysis results. Significant comparisons (p<0.05) shown in grey shading.

Stacked bar plots of the ten most abundant microbial families showed a dominance of *Enterobacteriaceae, Enterococcaceae* and, to a lesser extent, *Streptococcaceae* in libraries from diseased individuals compared to healthy ones (Figure 4). Differential expression analysis at the genus level identified 55 genera that were present in a significantly higher abundance in libraries comprising diseased individuals than healthy ones (padj <0.05, Supplementary Table 2). Genera were considered potential pathogens and analyzed further if they met the following criteria: (1) the genus was present in over half of the diseased libraries and (2) the genus was present in a significantly higher abundance in the diseased compared to the healthy samples in the differential expression analysis. Four bacterial genera met these criteria: *Streptococcus, Enterococcus, Corynebacterium* and *Escherichia.* However, as the genera *Corynebacterium* accounted for less than 1% of the total abundance in the diseased pools, it was excluded from further analysis. Within *Streptococcus, Enterococcus,* and *Escherichia*, contigs from 33 bacterial species were represented in our dataset. The dominant species within this subset were *Streptococcus gallolyticus*, *Enterococcus faecalis*, and *Escherichia coli,* accounting for 73% of the total abundance of these genera in metatranscriptomes from diseased individuals. These three species were subject to further analysis to investigate their potential association with exudative cloacitis.

**Figure 4.**
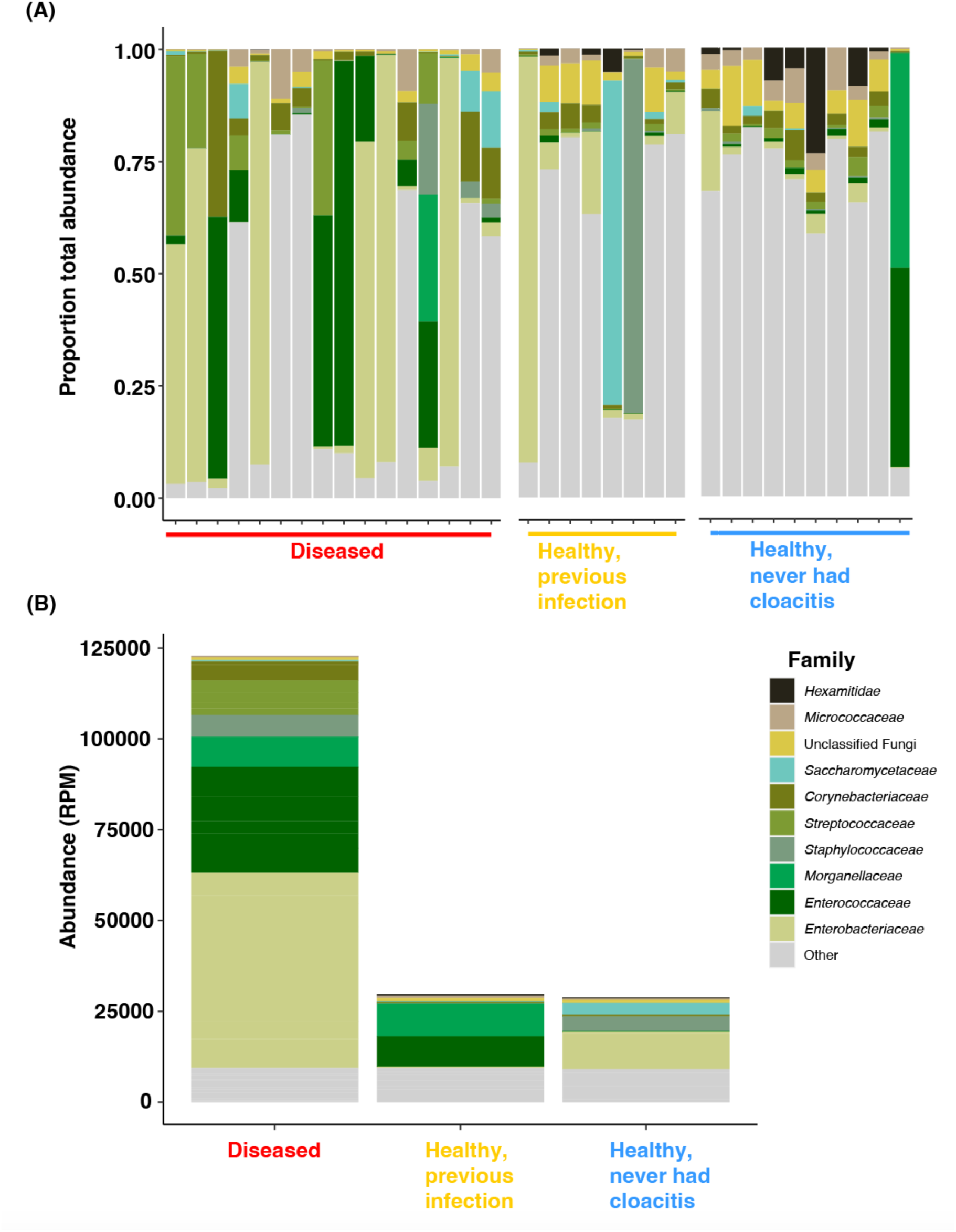
The proportional abundances per library (A) and total abundance in reads per million (B) of the ten most abundant microbial families. ‘Other’ refers to all other detected families not in the top ten most abundant.

For *S. gallolyticus* and *E. faecalis*, full MLST typing was often not possible with our short-read, RNA sequencing data as the individual alleles for each locus could not be matched together when multiple sequence types were present in the same library.

Instead, the abundance and phylogeny of each locus from the MLST scheme was analyzed individually.

### 3.4 S. gallolyticus

We were able to recover an average of four loci from each library out of the seven loci within the MLST scheme (Dumke et al., 2014). From libraries containing samples from healthy individuals, all extracted loci were novel alleles (i.e. they were not identical to any alleles in the PubMLST database). Phylogenetic analysis of each locus showed a strong clustering of these novel alleles, forming a divergent clade (Supplementary Figure 2). We also extracted alleles from a previous kākāpō shotgun metagenomics project (NETH00000000.1 (Waite et al., 2018)), and found these clustered with the alleles from our libraries of healthy kākāpō (Supplementary Figure 2). This may be evidence of a naturally present strain of *S. gallolyticus* that has evolved within kākāpō.

In contrast, five of the nine metatranscriptomes from diseased kākāpō had at least one allele that was an identical genetic match to alleles in the PubMLST database. These alleles were generally not consistent across these libraries (Supplementary Figure 2), suggesting multiple sequence types were present in the sampled kākāpō population. Samples from diseased kākāpō also had the same divergent, novel alleles as the healthy pools, and several had multiple alleles for each gene, suggesting multiple sequence types were present within some libraries, precluding full MLST typing for this species.

### 3.5 E. faecalis

We were not able to extract any of the seven MLST loci (Ruiz-Garbajosa et al., 2006) in the metatranscriptomic data from healthy kākāpō, suggesting *E. faecalis* was not present in these libraries. Within data from diseased kākāpō, we were able to extract an average of 4.5 loci per library out of the seven loci within the MLST scheme. Unlike *S. gallolyticus,* we only found one allele of each locus in each library, suggesting that only one sequence type was present in each library. However, these alleles were not consistent across libraries, and phylogenetic analysis showed they did not cluster together (Supplementary Figure 3). This indicates multiple sequence types were present in the sampled kākāpō population.

Three libraries had all seven loci from the MLST scheme, and only one allele from each locus was found in each library, allowing seven-gene sequence types (STs) to be inferred. Two of the three libraries (from two different individual kākāpō) were identified as containing ST287, while the other library contained ST63. ST287 has previously been found in diverse avian species in Europe (Aun et al., 2021; Jørgensen et al., 2017; Stępień-Pyśniak et al., 2018) as well as humans (hospitalized and non-hospitalized) in Europe, Ghana and Canada (PubMLST), while ST63 has previously been found in pigs (Shankar et al., 2006) and animal food in Europe and North America (PubMLST).

### 3.6 E. coli

We identified ten dominant *gnd* sequence types (gST) in our data (Figure 5A). The most abundant was gST171 (>5,800 RPM) followed by gST304 (3,500 RPM) and gST258 (3,000 RPM). As well as being the most abundant, gST171 was also found in the most libraries (13 libraries, 38%), followed by gST561 (12 libraries) and gST258 (10 libraries). The remaining seven most abundant *gnd* sequence types were each found in fewer than four libraries.

**Figure 5.**
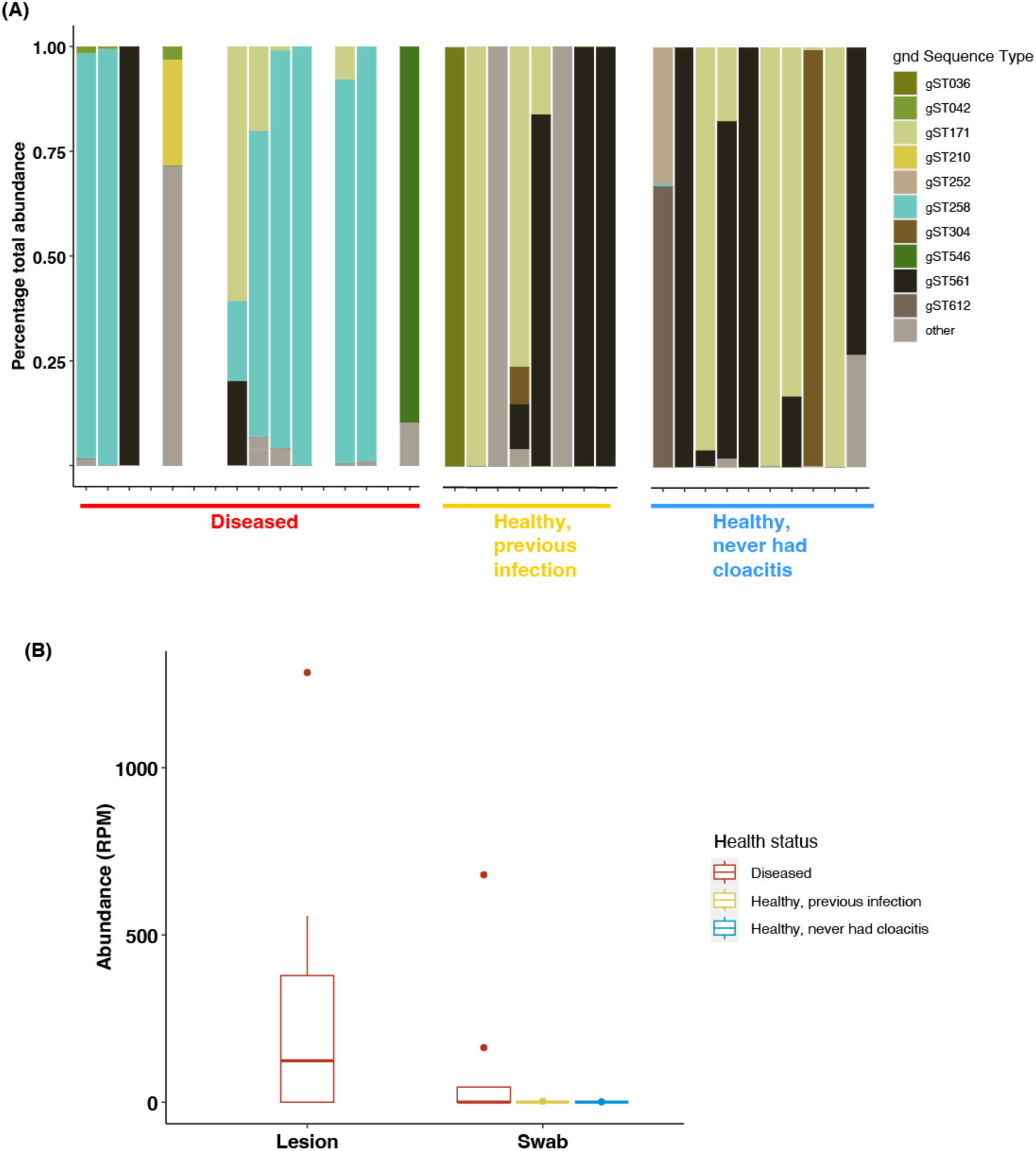
(A) The proportional abundances per library of the ten most abundant gnd sequence types, and (B) the abundance (reads per million, RPM) of gST258 in lesions and swabs from kākāpō of differing health status.

Metatranscriptomes from diseased kākāpō had a significantly higher abundance of gST258 than healthy kākāpō samples (p = 0.04, Supplementary Table 1, Figure 5B), and lesions had a significantly higher abundance than cloacal swabs (p = 0.01, Supplementary Table 1). The abundance of all other *gnd* sequence types were not significantly different in any comparisons. Phylogenetic analysis of the entire *gnd* gene for sequences belonging to gST258 showed the genes from kākāpō metatranscriptomes generally clustered within a large group from a variety of sample types including possum gut (New Zealand), sediment (New Zealand), human samples (USA, Czech Republic), wastewater (UK) and musk deer faeces (China) (Supplementary Figure 4).

A partial genome assembly was created from one data set from a diseased individual using a closely related whole genome as a reference (CP028770, isolated from possum faeces in Australia (Kang et al., 2021)). This reference was used because phylogenetic analysis of the *gnd* operon comparing our genome to other gST258 *E. coli* isolated from New Zealand and Australia indicated CP028770 was the most closely related annotated whole genome, with 99.9% nucleotide identity. The library was dominated by gST258 (accounting for 99.4% of *E. coli gnd* reads), and thus we could assemble a genome with reasonable accuracy. The assembled genome was 4.9 million base pairs (mbp) in length (including ambiguous bases), with a GC content of 51.2%. Coverage of the reference genome was 3.8 mbp (78.3%), with 93.2% identical bases. Other libraries had more variation in the sequence types present, and so we could not accurately assemble genomes. MLST analysis of this genome using the Achtman seven-locus MLST scheme showed that this genome belonged to ST141, and the O2/O50 O-antigen serogroup. The gene encoding the H antigen was not present in our partial genome assembly, thus the H antigen serotype could not be determined.

### 3.7 Virulence factors

Differential expression analysis identified 162 *S. gallolyticus* genes (out of a total of 2,171 genes in the reference genome) that were expressed at a significantly higher abundance in samples from diseased birds compared to the healthy ones (Supplementary Table 3). The key virulence genes that have been identified previously for *S. gallolyticus* are within the *pil1* pilus locus, consisting of collagen-binding domains, pilin motifs, and a LPXTG cell wall-anchoring domain (Danne et al., 2011). Of note, there was a highly significant difference in the expression of a gene producing the Cna B-type domain-containing protein (Figure 6, adjusted p-value padj = 0.000004), involved in collagen-binding (Danne et al., 2011). No pilin genes were significantly expressed, however for SpaH/EbpB family LPXTG-anchored major pilin this was driven by only one library from healthy kākāpō having expression of this gene. When this library was removed, the effect became significant (p = 0.03). D-alanyl-lipoteichoic acid biosynthesis proteins were also expressed at a significantly higher amount in diseased versus healthy kākāpō (Figure 6, *DltD* padj = 0.0005, *DltB*, padj = 0.006 and *DltA* padj = 0.046), which play a role in antimicrobial tolerance (Nilsson et al., 2016) and virulence (Du et al., 2023) in *Streptococcus mutans*. We also found a number of other genes significantly expressed in the diseased samples which are not necessarily associated with virulence, including the zinc ribbon domain containing protein (padj = 0.00004) involved in RNA polymerase function, YjjG family noncanonical pyrimidine nucleotidase (padj = 0.0000009) which breaks down mutagenic nucleotides (Titz et al., 2007), bifunctional lysozyme/C40 family peptidase (padj = 0.04) involved in bacterial cell wall degradation and remodelling, and tyrosine-type recombinase/integrase (padj = 0.01) involved in site-specific recombination and integration of DNA sequences (Li et al., 2020). We also found mannitol-1-phosphate 5-dehydrogenase (padj = 0.000009) and PTS mannitol specific transporter subunit IIBC (pdj = 0.00002) genes were highly expressed, both involved in sugar alcohol metabolism (Hu et al., 2020).

**Figure 6.**
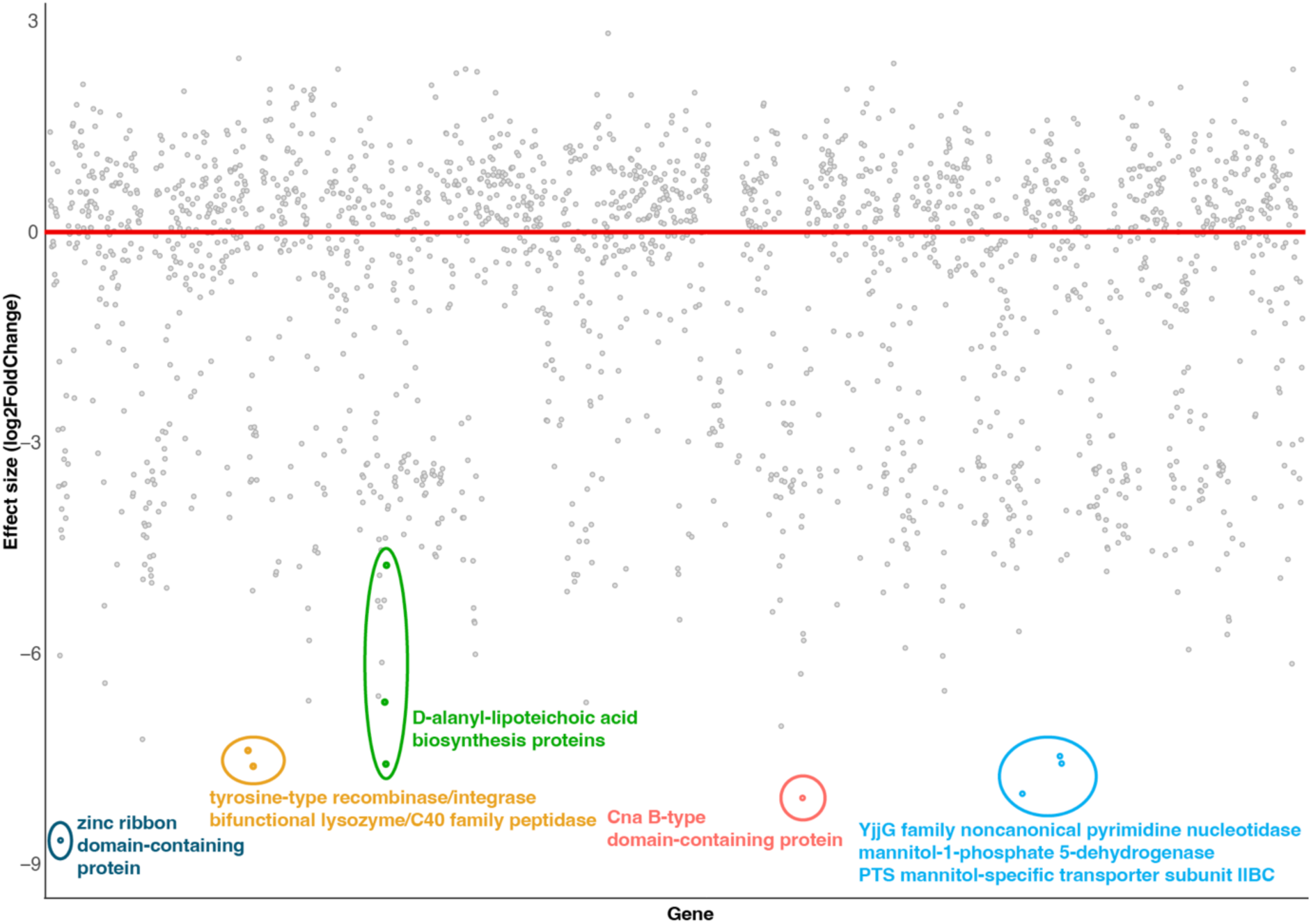
DiSerential expression analysis of Streptococcus gallolyticus genes (using NZ_CP054015.1 as the reference) for healthy vs diseased birds, showing overexpression of gene clusters in diseased samples, as shown by the coloured circles. The labels summarise the gene functions within each cluster. The red line shows an effect size of 0.

We identified 167 *E. coli* genes (out of a total 4,449 in the reference genome) that were expressed at a significantly higher abundance in the diseased kākāpō compared to the healthy ones (Supplementary Table 4). All nineteen genes from the colibactin biosynthesis gene cluster were expressed at a significantly higher abundance in the diseased compared to the healthy metatranscriptomes (*ClbA – S*, Supplementary Table 4, Figure 7). This gene cluster synthesizes colibactin, which damages DNA in host cells (Vizcaino & Crawford, 2015). Genes in the salmochelin system involved in iron acquisition (Gao et al., 2012) were also highly expressed in samples from diseased kākāpō, with all five genes in the system expressed at a significantly higher abundance in the diseased than the healthy kākāpō (Figure 7, siderophore salmochelin receptor *IroN* padj = 0.001, catecholate siderophore esterase *IroE* padj = 0.01, salmochelin/enterobactin export ABC transporter *IroC* padj = 0.01, salmochelin biosynthesis C-glycosyltransferase *IroB* padj = remod0.003). Both colibactin and the salmochelin system are associated with virulence in extraintestinal pathogenic *E. coli* (ExPEC) in birds (Gao et al., 2012; Johnson et al., 2008). Also of note, several genes associated with adhesion and promoting persistence in uropathogenic *E. coli* (UPEC, a subset of ExPEC) (Figure 7, the ag43 gene p = 0.0004, and S fimbrial adhesion genes *SfaE-G*, *SfaX* and *SfaY*) were significantly highly expressed in the diseased compared to the healthy kākāpō (Klemm et al., 2010; Lüthje & Brauner, 2010). Two multidrug/spermidine efflux SMR transporter subunits were also significantly expressed (*MdtJ* p = 0.0005, *MdtI* p = 0.047), involved in multidrug resistance (Burata et al., 2022). A cluster of genes involved in lipid metabolism and transport were also highly expressed, (e.g., beta-ketoacyl-ACP synthase, acyl carrier protein, acyl-CoA synthetase), as well as the type VI secretion system which plays a role in attacking eukaryotic host cells and competing against other bacteria (Lien & Lai, 2017). Genes involved in genetic mobility (primarily transposase genes) were also highly expressed across multiple gene clusters (Figure 7).

**Figure 7.**
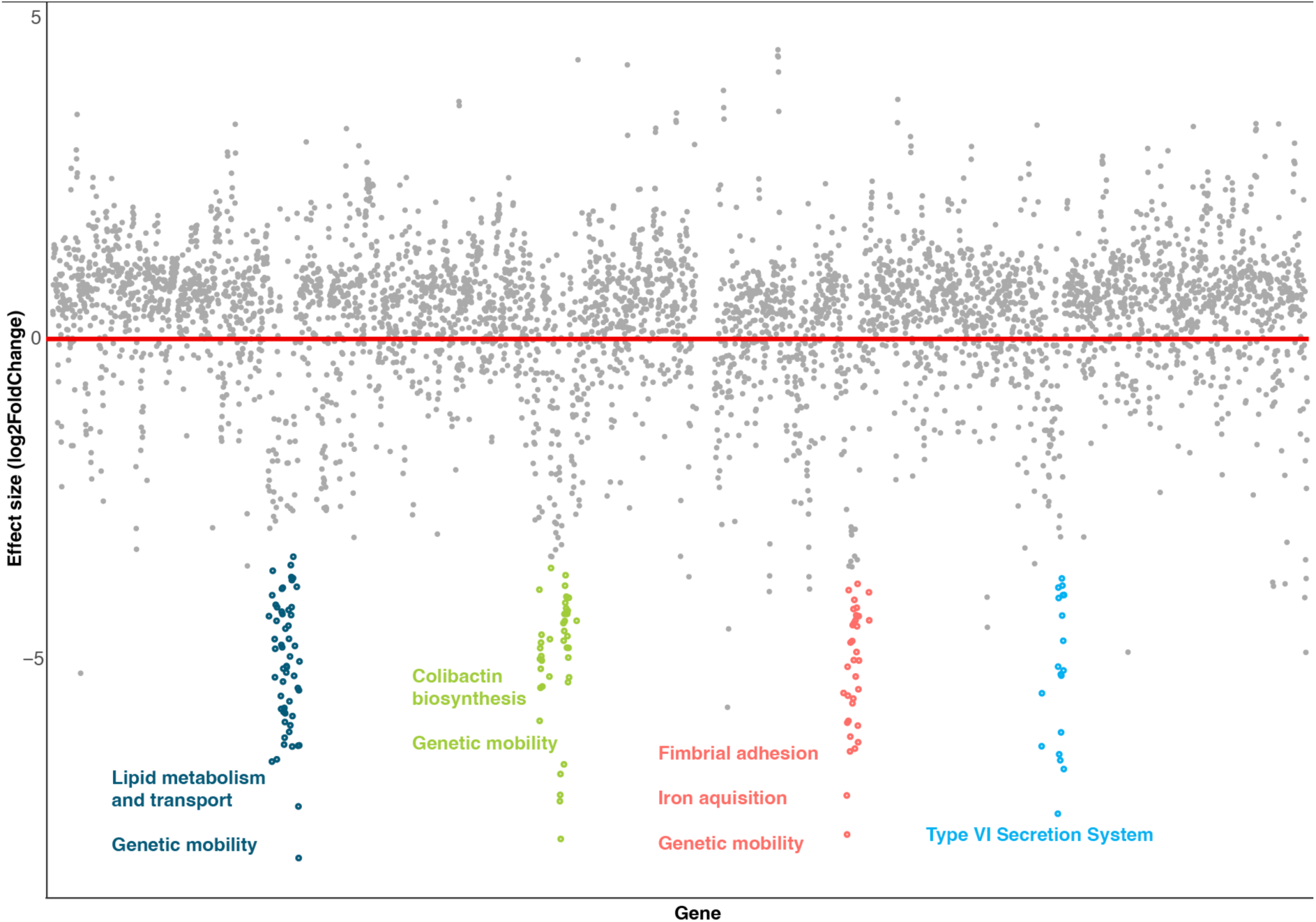
DiSerential expression analysis of Escherichia coli genes (using CP028770 as the reference) for healthy vs diseased birds, showing overexpression of gene clusters in diseased samples, as shown by the coloured circles. The labels summarise the gene functions within each cluster. The red line shows an effect size of 0.

As *E. faecalis* did not appear to be present in metatranscriptomes from healthy kākāpō, we did not compare individual gene expression levels between samples from diseased and healthy individuals. Within those from diseased birds, the most abundantly expressed gene across the entire genome (3,282 genes in the reference genome) was the virulence gene pilus tip protein *EbpA*, considered an important contributor to the pathogenicity of *E. faecalis* (Nielsen et al., 2012) (Figure 8, Supplementary Table 5).

**Figure 8.**
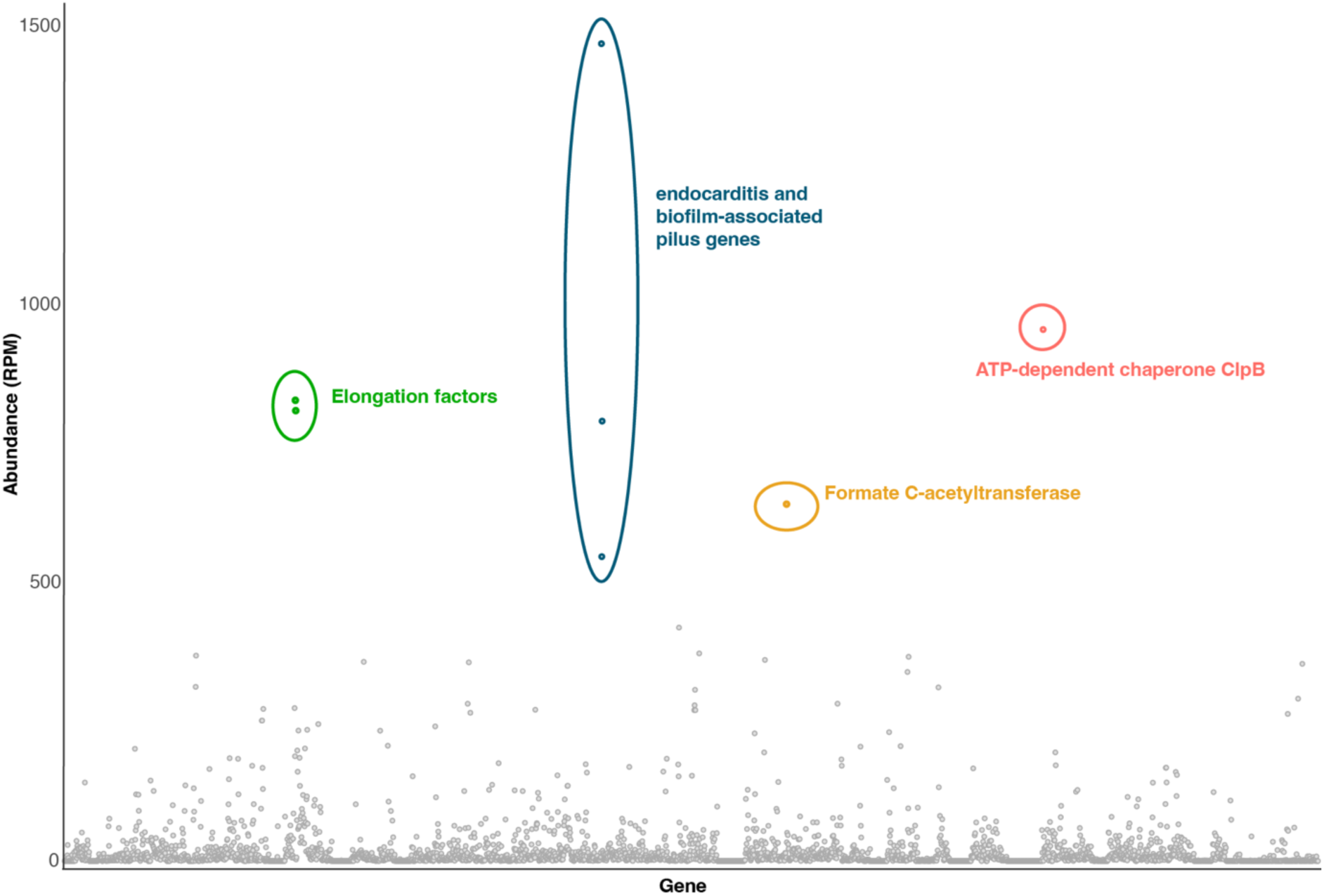
Expression of Enterococcus faecalis genes (using NZ_KE136528.1 as the reference) in diseased samples, expressed as reads per million (RPM). The colours and labels denote the highly expressed genes.

Transcripts for this gene were present at 1,468 RPM, present in 10 out of the 16 libraries from diseased individuals, and accounted for 2% of the total reads aligned to the genome. The other genes that make up the pili were also expressed highly, with the pilus major subunit *EbpC* the fifth most abundantly expressed gene (790 RPM, 10 libraries), and the pilus minor subunit *EbpB* the seventh (547 RPM, 12 libraries). Also of note, we found ATP-dependent chaperone *ClpB* was highly expressed (955 RPM, 12 libraries), which plays an important role in virulence and antimicrobial resistance in *E. faecalis* (Zheng et al., 2020). We also found the formate C-acetyltransferase gene (involved in glucose metabolism) was highly expressed, along with elongation factors *G* and *Tu* involved in protein synthesis (Andersen et al., 2003).

### 3.8 Antimicrobial resistome

We detected 16 antimicrobial resistance (AMR) genes that had an abundance (i.e. gene expression) of >1 RPM. However, fourteen of these were detected in only one library each, suggesting they were present at a low frequency in the kākāpō population. The remaining two AMR genes (*mupA* and *lsa(A)*) were present at >1RPM in five libraries each. *MupA* was present in one library from a diseased individual and four libraries from healthy individuals. *lsa(A)* was present in five libraries from diseased individuals and no libraries from healthy individuals. Importantly, the *lsa(A)* gene is an intrinsic gene in *E. faecalis* (Singh et al., 2002), which explains why the gene was not present in libraries from healthy birds. Both genes were present at relatively low abundances, with the highest being 11 RPM (*mupA*) and 67 RPM (*lsa(A)*). *lsa(A)* was significantly more abundant in the lesions than the swabs (p = 0.02, Supplementary Table 1).

We also detected two AMR point mutations (*gyrA* and *parC*) for *E. faecalis* at a significantly higher abundance in the lesions than the swab libraries (p = 0.009, p = 0.002 respectively, Supplementary Table 1). *parC* was also significantly more abundant in the diseased than healthy libraries (p = 0.03), while *gyrA* was not (p = 0.1, Supplementary Table 1).

We detected thirteen AMR point mutations for *E. coli*, at relatively high abundances (average 394 RPM, range 0 – 7518). However, none of these genes had significant differences in abundance between sample type or health status.

## 4. Discussion

Our study investigated the microbial communities associated with exudative cloacitis in the critically endangered kākāpō, uncovering potential pathogens, their virulence factors, and antimicrobial resistance gene expression. These findings shed light on possible causative agents and offer insights into the interplay of microbial species that could be influencing the disease.

We identified three bacterial species – *S. gallolyticus*, *E. faecalis*, and *E. coli* – that were significantly more abundant in diseased kākāpō compared to healthy individuals. Notably, the genera that these bacterial species belong to – *Streptococcus, Enterococcus* and *Escherichia* respectively – have been shown to be differentially expressed in cloacitis cases previously (West et al., 2024), lending weight to the hypothesis that these bacteria are linked to the disease. These three bacterial species are all known pathogens, both in avian (Chadfield et al., 2004; Hess et al., 2021; McPeake et al., 2005) and human hosts (Danne et al., 2011; Gao et al., 2012; Nielsen et al., 2012), however they are also frequently found in diverse environmental settings, including soil and water (Hammerum, 2012; Xing et al., 2019), as well as commensally in the gastrointestinal tracts of birds (Cagri et al., 2024; McPeake et al., 2005; Stępień-Pyśniak et al., 2019). Therefore, the presence of virulence factors is an important indication of whether a particular bacteria is commensal or pathogenic (Keen, 2012).

We found *S. gallolyticus* displayed elevated expression of a gene expressing the Cna B-type domain-containing protein, a virulence factor involved in collagen binding and biofilm formation, which has been implicated in endocarditis in humans (Danne et al., 2011). These results suggest *S. gallolyticus* may be contributing to exudative cloacitis in kākāpō. Similarly, *E. faecalis* genes related to biofilm production and endocarditis, such as the *EbpA, EbpB*, and *EbpC* pilus proteins were highly expressed, suggesting a role for biofilm formation in disease progression (Nielsen et al., 2012). Importantly from a conservation perspective, the genetic diversity observed in both *S. gallolyticus* and *E. faecalis* among diseased kākāpō suggests that these bacteria originate from exogenous sources rather than from kākāpō-to-kākāpō transmission. This source could be environmental such as soil or water, or cross-species transmission from other species. These findings indicate the need for further investigation into the environment on the islands with kākāpō populations, to determine if these bacteria are present and to identify potential sources of infection.

In *E. coli*, we found overexpression of genes associated with iron acquisition and the colibactin genotoxin, both of which play key roles in bacterial virulence. The salmochelin siderophore system enables *E. coli* to scavenge iron under iron-limited conditions within the host, enhancing its survival and virulence during infection (Gao et al., 2012). Additionally, the colibactin biosynthesis gene cluster was highly expressed, suggesting the production of this genotoxin, which induces DNA damage in host cells and may contribute to immune evasion and pathogenicity (de Oliveira Alves et al., 2024; Vizcaino & Crawford, 2015). The presence of colibactin is particularly significant as epidemiological studies have shown that the colibactin biosynthesis gene cluster was significantly associated with a highly virulent subset of extraintestinal pathogenic *E. coli* (ExPEC) (Johnson et al., 2008).

The interplay between the salmochelin siderophore system and the colibactin biosynthesis pathway in *E. coli* is also important, particularly in terms of their combined contribution to bacterial virulence in relation to iron (Martin et al., 2017). Colibactin synthesis is closely regulated by iron bioavailability (Martin et al., 2017). Thus, the salmochelin system not only enhances *E. coli’*s ability to thrive in the host by providing essential iron but also supports the production of colibactin by ensuring the bacteria have the necessary resources to synthesize the genotoxin (Martin et al., 2017).

Furthermore, research in poultry with colibacillosis revealed the presence of *E. faecalis* further augments avian pathogenic *E. coli* (APEC) survival and growth under iron limiting conditions, possibly translating to the increased virulence of APEC (Walker et al., 2020). A similar interaction could be occurring in kākāpō, but further work is needed to confirm this. Unlike West et al. (2024), we did not find an association with disease and the expression of the toxin producing *hok*/*gef* genes. This difference may be due to the sample types: West et al. (2024) found elevated expression of these genes primarily in faecal samples, whereas our study focused on cloacal swabs and lesion samples, which might more accurately represent the microbiome specific to the cloacal environment.

The identification of *E. coli* ST141 in our dataset is notable as this sequence type belongs to phylotype B2 in the extraintestinal pathogenic *E. coli* (ExPEC) group (Emery et al., 2023), and uropathogenic *E. coli* (UPEC) that cause urinary tract infections in humans (Gati et al., 2019). ExPEC strains are known for causing infections outside the intestinal tract in both humans and animals, including avian species (Mellata, 2013; Smith et al., 2007). The presence of ExPEC-associated virulence factors in diseased kākāpō further suggests that *E. coli* may play a critical role in disease progression by facilitating iron acquisition and causing DNA damage in host cells. These results, combined with the consistent presence of one *E. coli gnd* sequence type across diseased birds suggests that this may be the primary pathogen causing this disease (perhaps in conjunction with *E. faecalis*), however more study is needed to confirm this. Interestingly, phylotype B2 is associated with ‘low impact’ exotic or natural forest sites elsewhere in New Zealand (Cookson et al., 2024; Cookson et al., 2022), possibly suggesting the source of this bacteria is not human associated, although this may not be the case for the *E. coli* we identified in kākāpō.

Despite thorough analysis, including both sequence and structural similarity searches, we did not identify any viruses directly infecting kākāpō. While we detected a diverse virome, the majority of viral sequences were plant and bacteria host-related, with no evidence of kākāpō-specific viruses. This finding suggests that viral infection is unlikely to be the primary cause of exudative cloacitis. Another possibility is that the primary pathogen of this disease is in fact a virus, but it is cleared quickly and thus not detected by our sampling. Viral infection could predispose kākāpō to secondary bacterial disease. Indeed, *Infectious bronchitis virus* (IBV) predisposes chickens to colibacillosis caused by avian pathogenic *E. coli* (Ariaans et al., 2008). As kākāpō are diagnosed and sampled following capture in the wild, date of disease onset is unknown and sampling likely occurs weeks or months after disease onset.

Given that antibiotics are the primary treatment for cloacitis in kākāpō, there are concerns that frequent antibiotic use could drive the emergence of AMR in these birds. We observed the presence of several AMR genes in diseased kākāpō. Mutations in the *gyrA* and *parC* genes were detected, both of which are key components of the quinolone resistance-determining regions (QRDR) (Kim & Woo, 2017). The presence of these mutations suggests that fluoroquinolones, used in veterinary medicine (Goggs et al., 2021) and commonly used in kākāpō until recently (unpublished data), may be less effective in treating infections caused by these resistant bacteria. We also identified two multidrug/spermidine efflux SMR transporter subunits involved in multidrug resistance (Burata et al., 2022). These findings highlight the significant challenge of AMR in the management of bacterial infections in endangered species like kākāpō, where treatment options are already limited.

Pathogen discovery is challenging, particularly in wildlife. One challenge, as discussed above, is the timing of sample collection as many infected animals are likely diagnosed, sampled and treated long after initial disease onset. Additionally, the inability to conduct experimental challenge studies in endangered species limits our ability to confirm causality. In kākāpō, conducting experimental infections to reproduce the disease is ethically and logistically impossible due to the risk of further harming a critically endangered population. This means we cannot definitively prove that a particular microorganism is the causative agent of exudative cloacitis or other diseases. This limitation means that alternative methods, such as observational studies and molecular analysis, are relied upon to infer causality, but these approaches cannot provide the same level of certainty as experimental challenge studies.

Despite these challenges, RNA sequencing is an invaluable tool for identifying potential pathogens associated with disease, particularly in endangered species where the number and type of samples is limited. The ability to uncover the transcriptomes from a wide array of infectious agents, including bacteria, fungi, and viruses, provides a comprehensive view of the microbial ecosystem associated with disease. Furthermore, the technique’s unbiased approach allows for the detection of novel or unexpected pathogens, which may have been missed using more targeted methods. However, the sheer diversity of RNA present in each sample creates challenges in sequencing depth and specificity, particularly when attempting to distinguish between closely related bacterial strains. In our study, the limitations of metatranscriptomics prevented us from definitively classifying some bacterial strains using MLST schemes. Despite these challenges, this approach provides a broad, unbiased view of the infectome, enabling the identification of microorganisms across bacterial, viral, and fungal domains. In this study, it has been especially useful for identifying potential bacterial pathogens and their virulence mechanisms, guiding future conservation and disease management strategies.

By identifying the specific pathogens and virulence factors associated with cloacitis in kākāpō, our results lay the groundwork for practical conservation interventions and identifies key areas for further research. First, understanding the microbial agents responsible for the disease can inform targeted treatment approaches, potentially allowing for more effective and specific antibiotic or antimicrobial therapies that minimize the risk of resistance (Harrison et al., 2020). Research pinpointing environmental sources or carriers of these pathogens could enable proactive management strategies, such as habitat modification and biosecurity measures (Shapiro et al., 2021). Lastly, the identification of key virulence factors could facilitate the development of vaccines or immune-boosting strategies tailored to the kākāpō’s unique physiology, providing a long-term tool to mitigate the impact of cloacitis (Wu et al., 2008). Together, these insights contribute to a holistic approach that combines immediate treatment options with preventative measures, ultimately supporting the survival and health of this critically endangered species.

## Supplementary Table and Figure legends

**Supplementary Figure 1.** Ormycovirus phylogeny and conserved motifs. Maximum likelihood midpoint rooted phylogenetic tree (a.) of representative ormycovirus transcripts containing the RdRp. The kākāpō associated ormycovirus identified in this study is bolded while viruses that were identified through screening the TSA are noted. Branches are scaled to the number of amino acid substritutions per site. Nodes with ultrafast bootstrap values of >90% are noted by an asterisk. Below the phylogeny is an alignment of ormycovirus RdRp amino acid sequences (b.). Conserved motifs (A-C) are shown while the kākāpō associated ormycovirus is bolded.

**Supplementary Figure 2.** Maximum likelihood phylogenetic tree of the seven *Streptococcus gallolyticus* loci used in multi-locus sequence typing (MLST). Sequences found in healthy libraries are shown as blue labels, sequences found in diseased libraries shown in red. Alleles from the PubMLST database are shown in black. The sequences extracted from a previous kākāpō shotgun metagenomic project (Waite et al., 2018), are also shown in black (NETH01000002.1 - NETH01000038.1). Branches are scaled according to the number of amino acid substitutions per site, shown in the scale bars. The trees are midpoint rooted for display purposes only. Black circles on nodes show bootstrap support values of more than 85%.

**Supplementary Figure 3.** Maximum likelihood phylogenetic tree of the seven *Enterococcus faecalis* loci used in multi-locus sequence typing (MLST). Sequences found in diseased libraries shown in red. Alleles from the PubMLST database are shown in black. Black circles on nodes show bootstrap support values of more than 85%.

**Supplementary Figure 4.** Maximum likelihood phylogenetic tree of the *Escherichia coli gnd* gene. Sequences found in diseased libraries shown in red, related sequences are shown in black. Branches are scaled according to the number of amino acid substitutions per site, shown in the scale bar.

**Supplementary Table 1**. Results of linear, linear mixed effects, generalised linear mixed effects and PERMANOVA models comparing sample types and disease status.

**Supplementary Table 2**. DESeq2 output for the infectome dataset, comparing diseased versus healthy samples.

**Supplementary Table 3**. DESeq2 output for the *Streptococcus gallolyticus* gene dataset, comparing diseased versus healthy samples.

**Supplementary Table 4**. DESeq2 output for the *Escherichia coli* gene dataset, comparing diseased versus healthy samples.

**Supplementary Table 5**. Rsubread output for the *Enterococcus faecalis* gene dataset, showing the total abundance (in reads per million) of each gene and the number of libraries it was detected in.

## Acknowledgements

Thank you to Dr Simon Jackson and Dr Leighton Payne for their advice on protein structural analyses. The authors wish to acknowledge the Morrison Foundation for their support to the Kākāpō Recovery Team. The authors used the New Zealand eScience Infrastructure (NeSI) high performance computing facilities as part of this research.

New Zealand’s national facilities are provided by NeSI and funded jointly by NeSI’s collaborator institutions and through the Ministry of Business, Innovation & Employment’s Research Infrastructure programme. URL https://www.nesi.org.nz.

## Data Availability

Sequence data generated in this study has been deposited in the Sequence Read Archive (SRA) under the accession number XXXX.

## Ethics

The data used in this study were collected as part of routine kākāpō conservation management conducted by the New Zealand Department of Conservation (NZDOC) as required by the New Zealand Conservation Act (1987). This study was exempt from the requirement of animal ethics approval under NZDOC’s obligations to the New Zealand Animal Welfare Act (1999).

