## Supplementary figures and images for "Evidence for a role of extraintestinal pathogenic *Escherichia coli, Enterococcus faecalis* and *Streptococcus gallolyticus* in the aetiology of exudative cloacitis in the critically endangered kākāpō (*Strigops habroptilus*)"

### supplementary_figure1_TSA_alignment_trimmed.fasta.tree.pdf

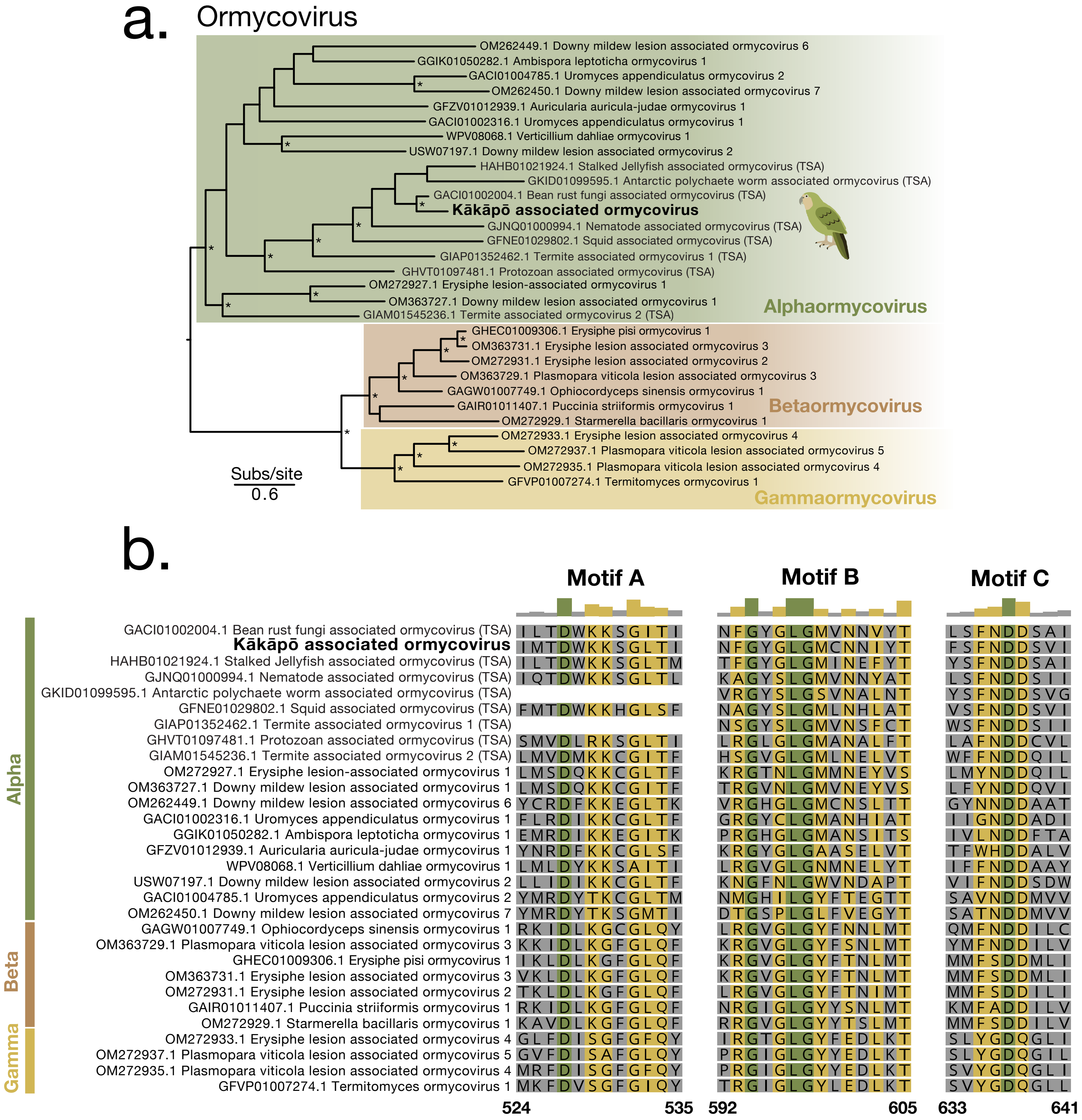

### supplementary_figure2_Strepgall_trees.pdf

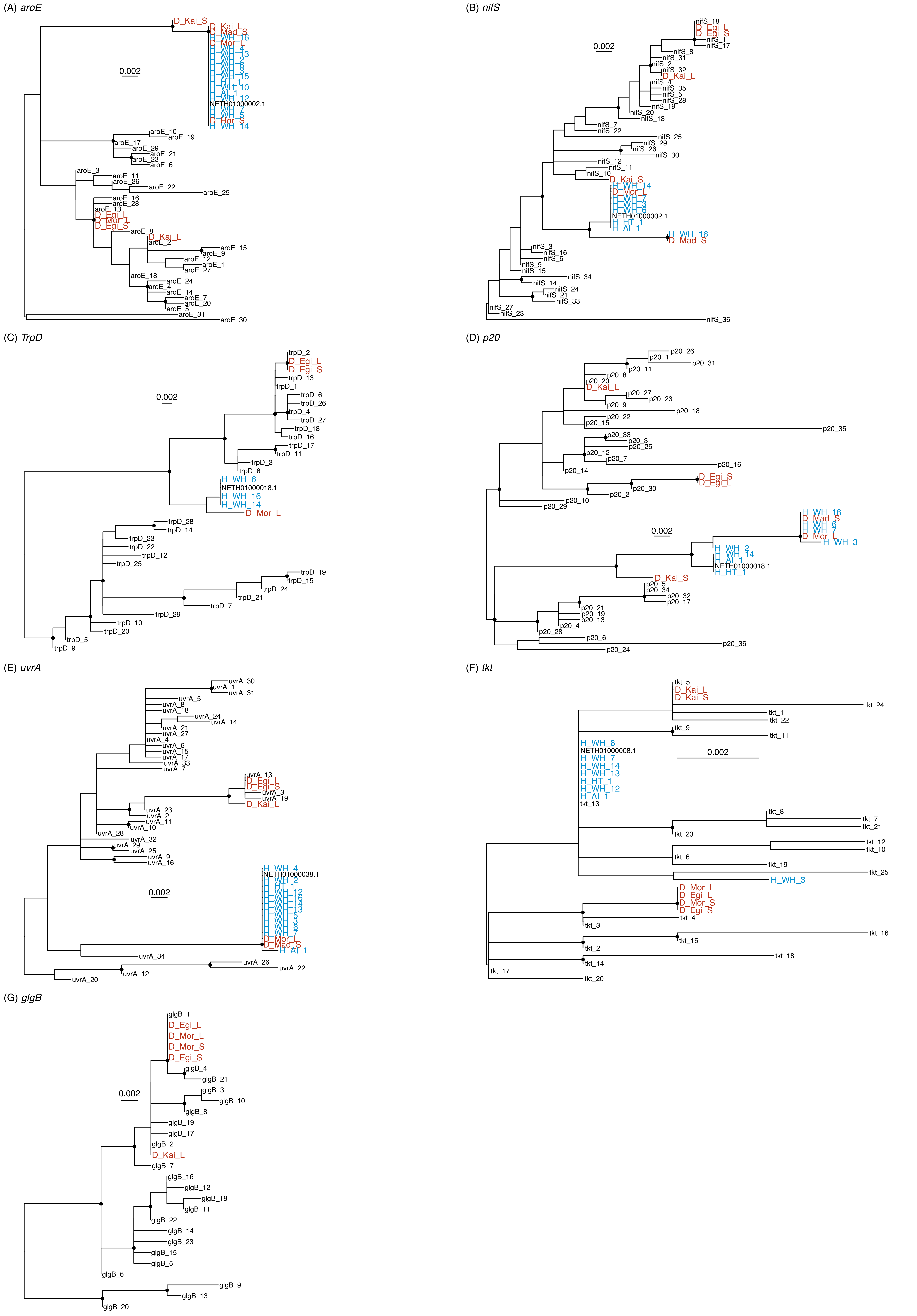

### supplementary_figure3_Efaecalis_trees.pdf

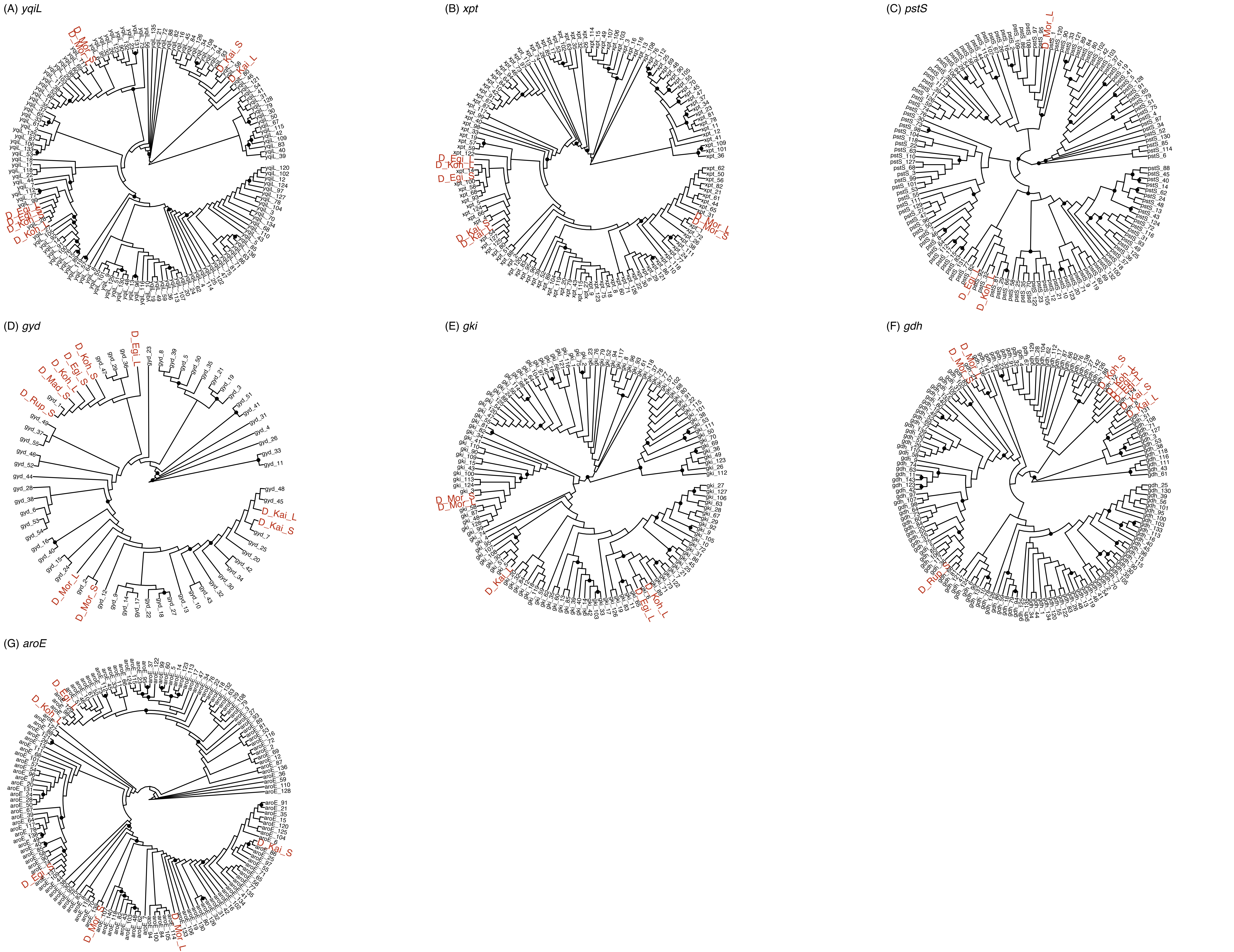
